## Supplementary figures and images for "Dual specificity phosphatase 7 drives the formation of cardiac mesoderm in mouse embryonic stem cells"

### Supplementary Figure 1a

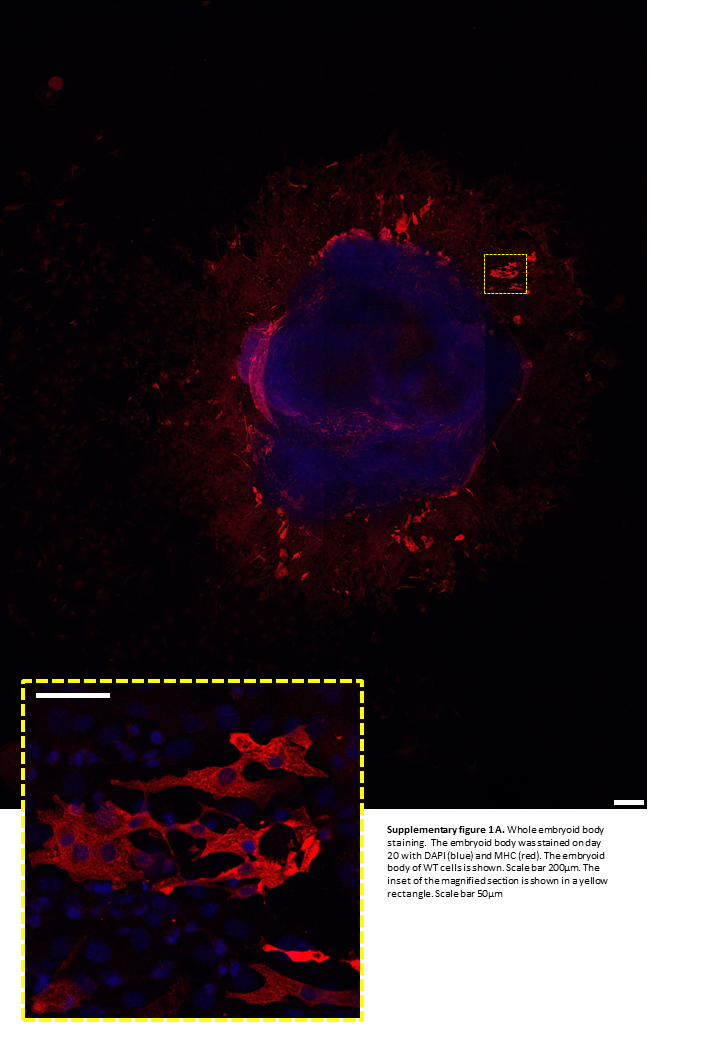

### Supplementary Figure 1c

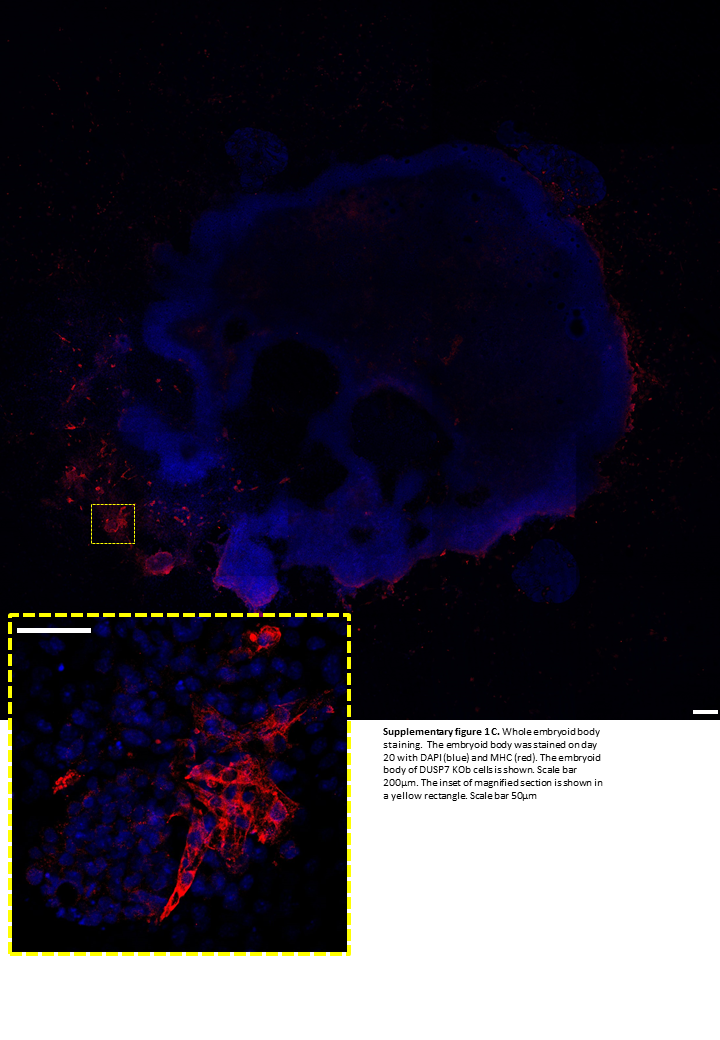

### Supplementary Figure 1d

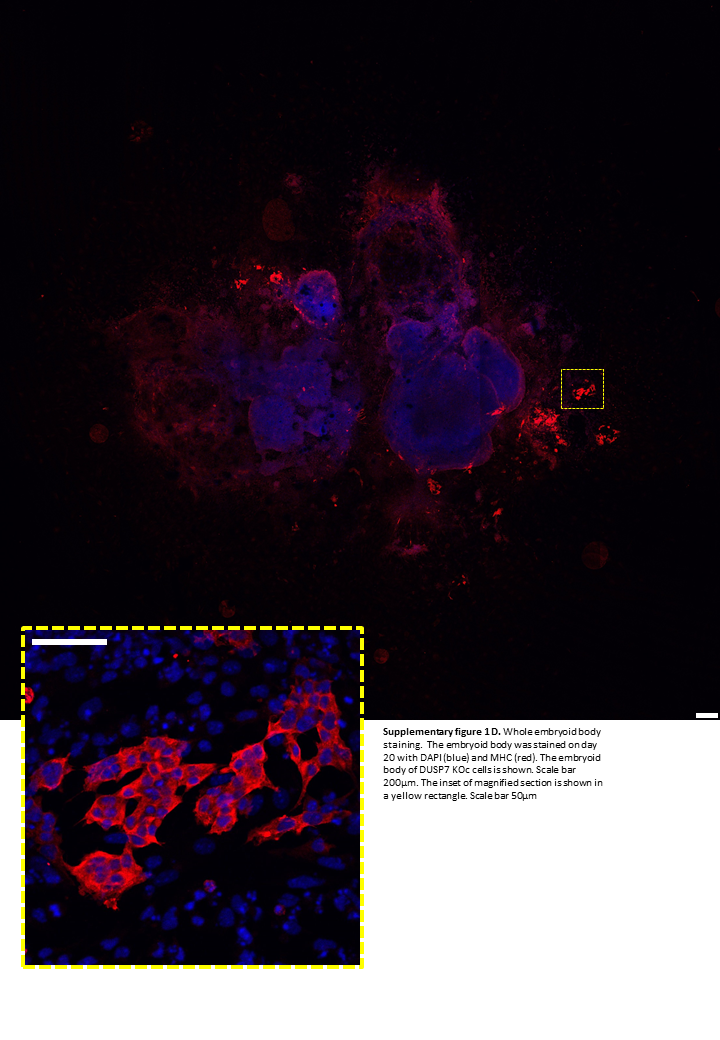

### Supplementary Figure 2a

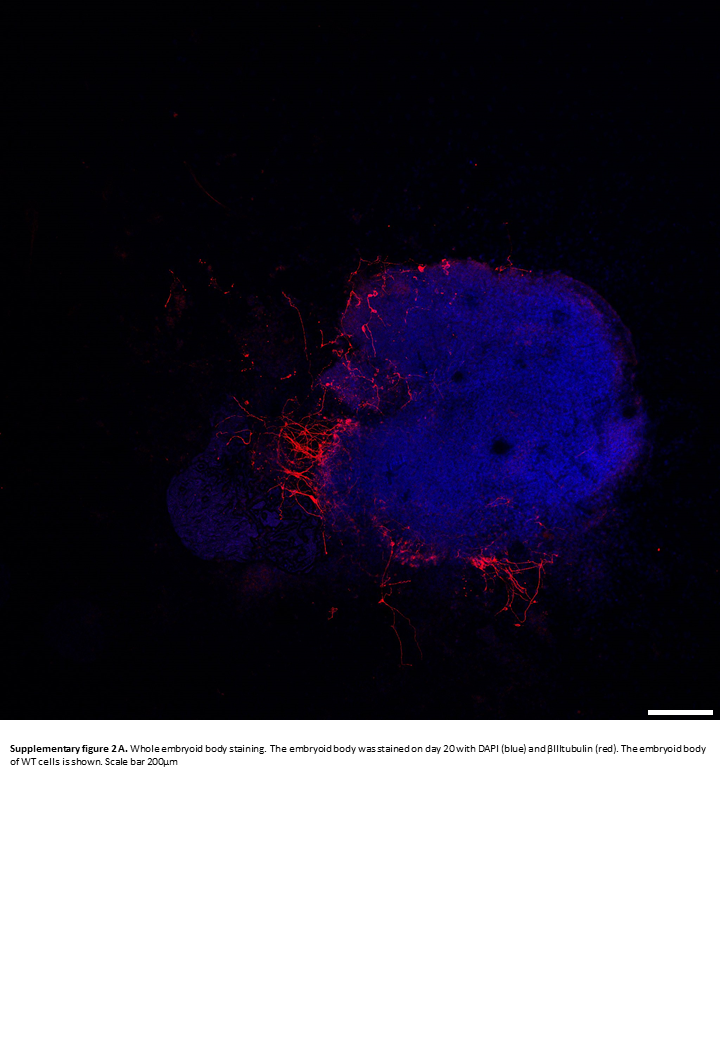

### Supplementary Figure 2b

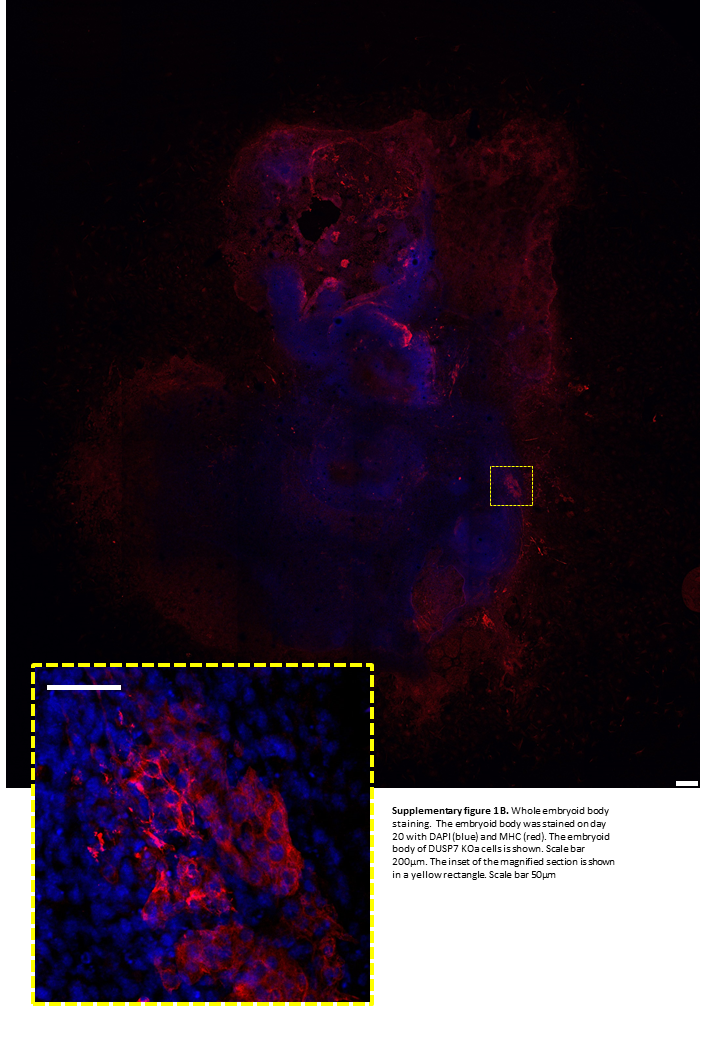

### Supplementary Figure 2b

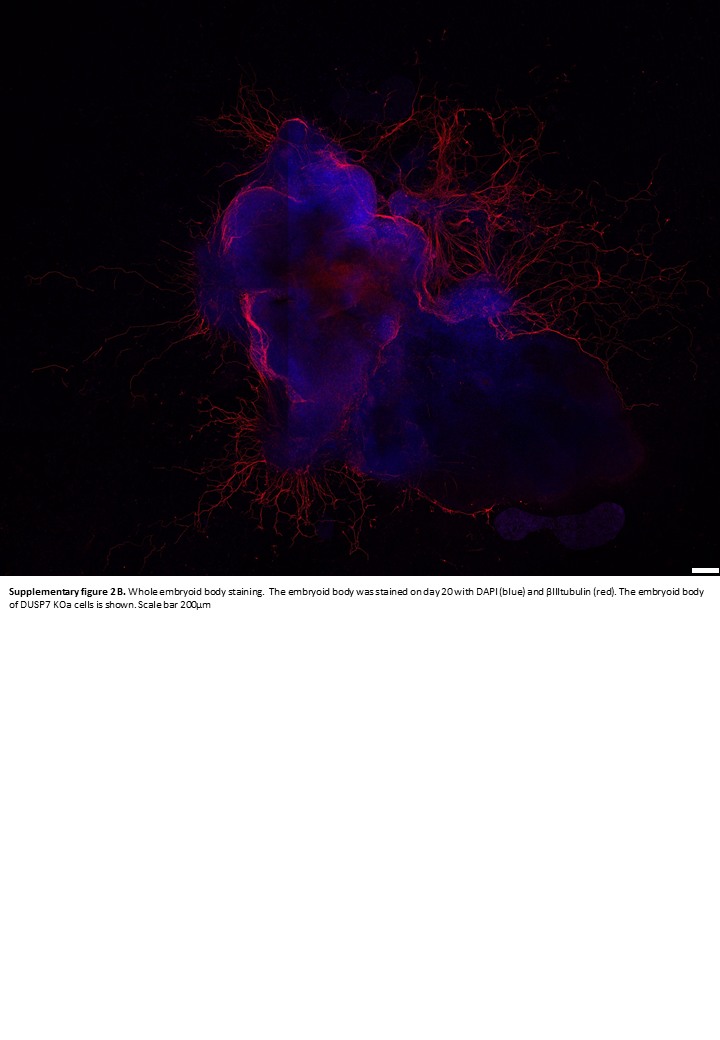

### Supplementary Figure 2c

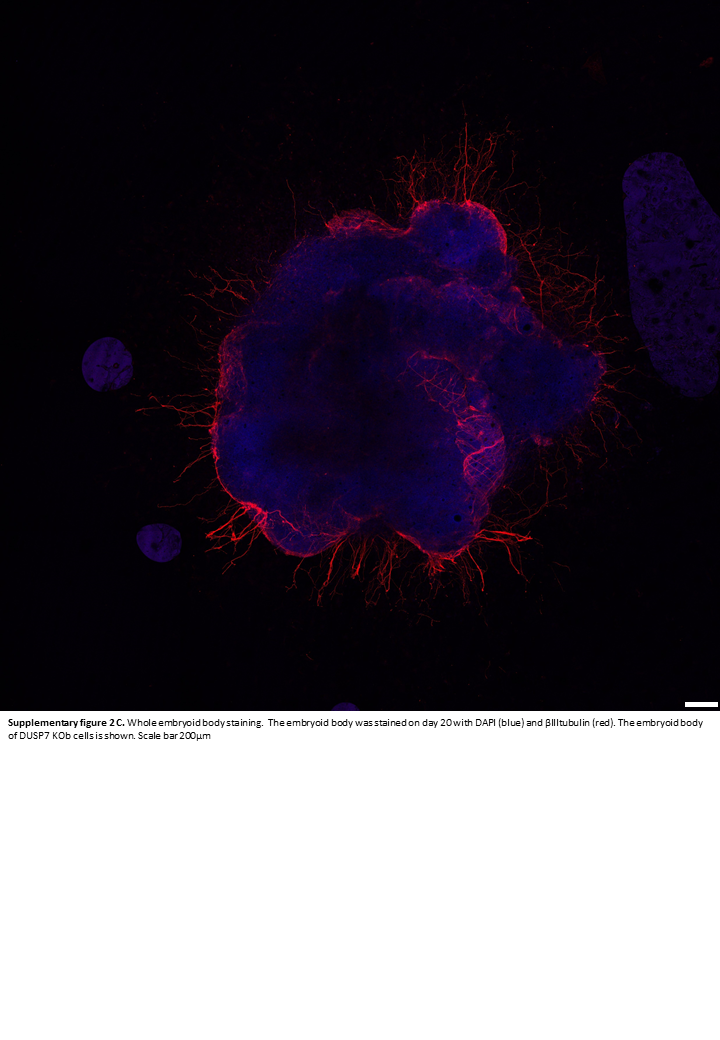

### Supplementary Figure 2d

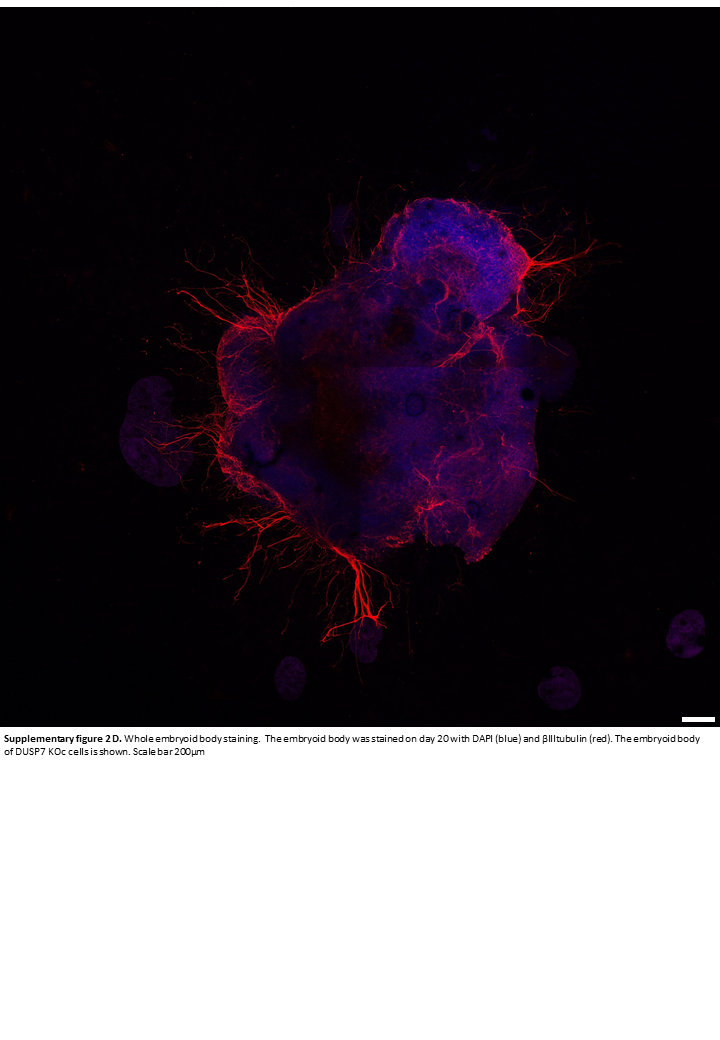

### Supplementary Figure 3a

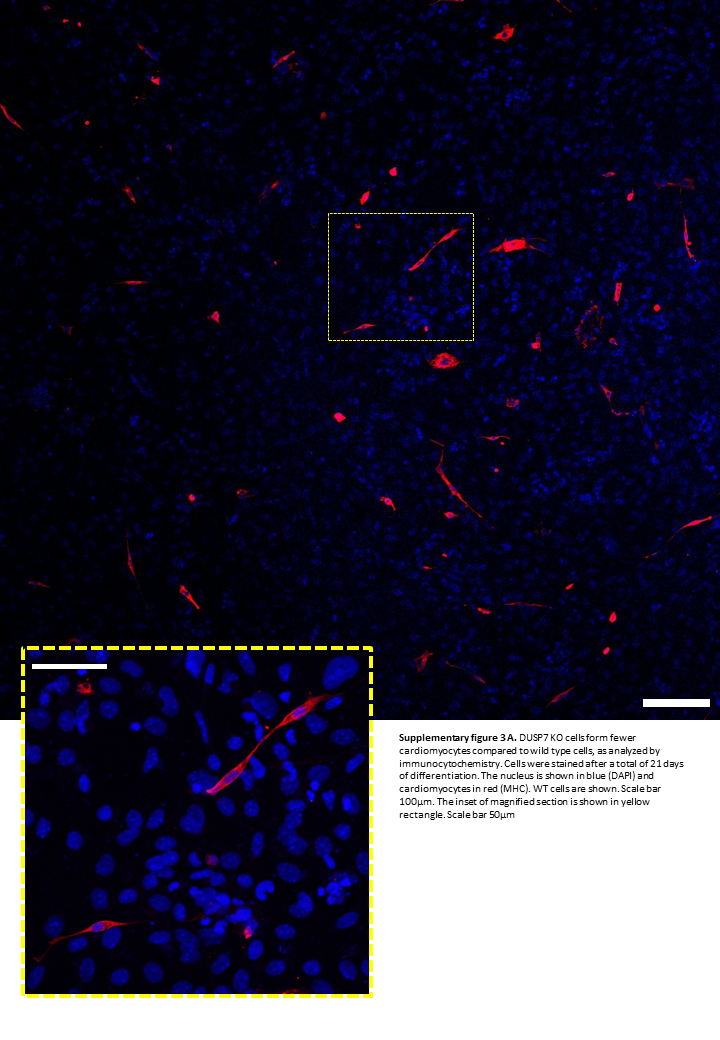

### Supplementary Figure 3b

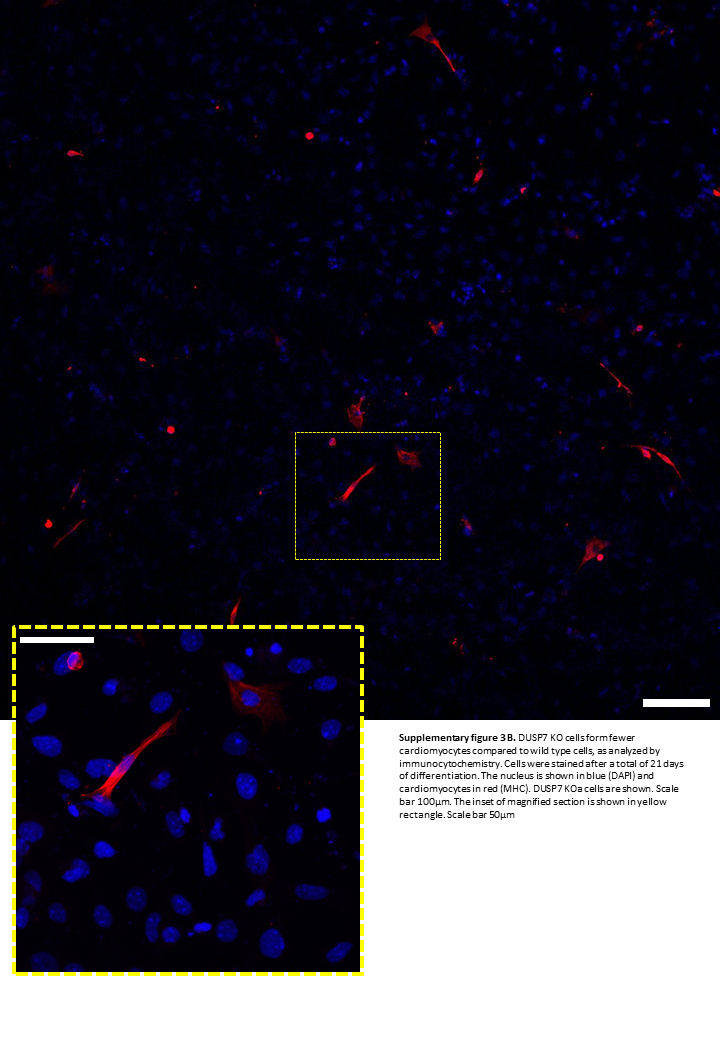

### Supplementary Figure 3c

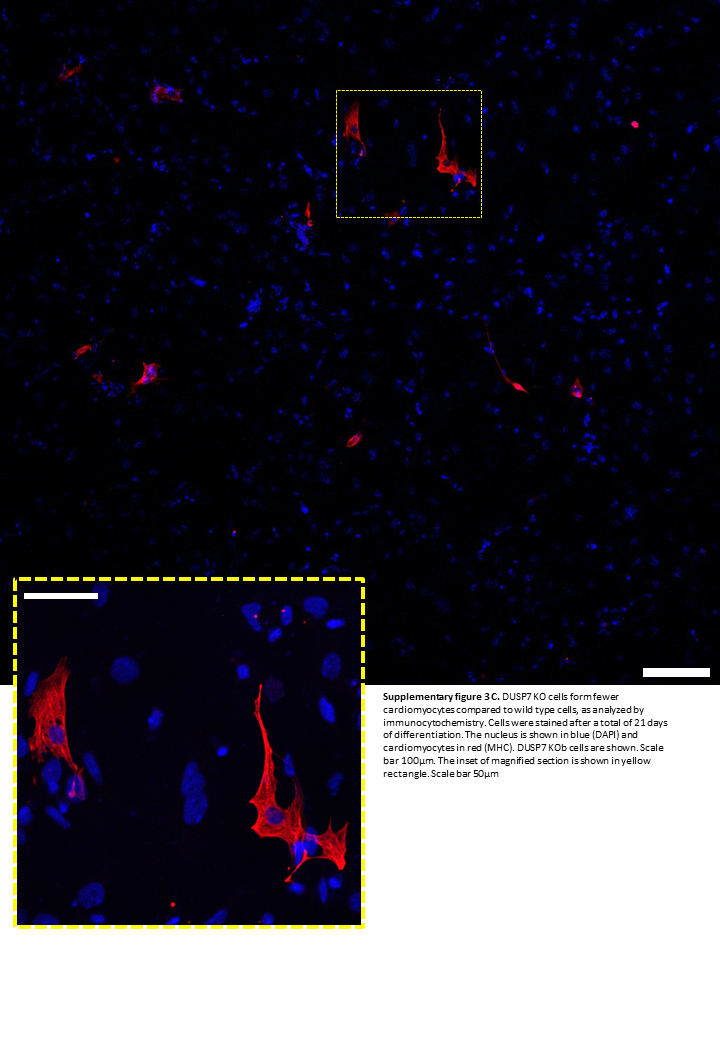

### Supplementary Figure 3d

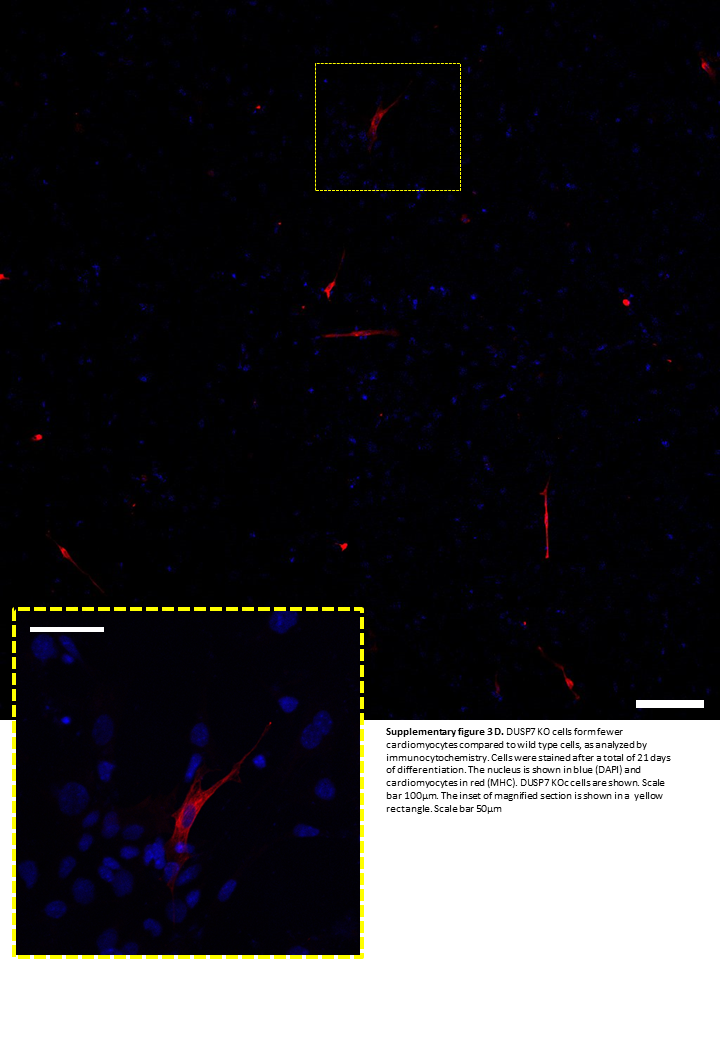

### Supplementary Figure 4

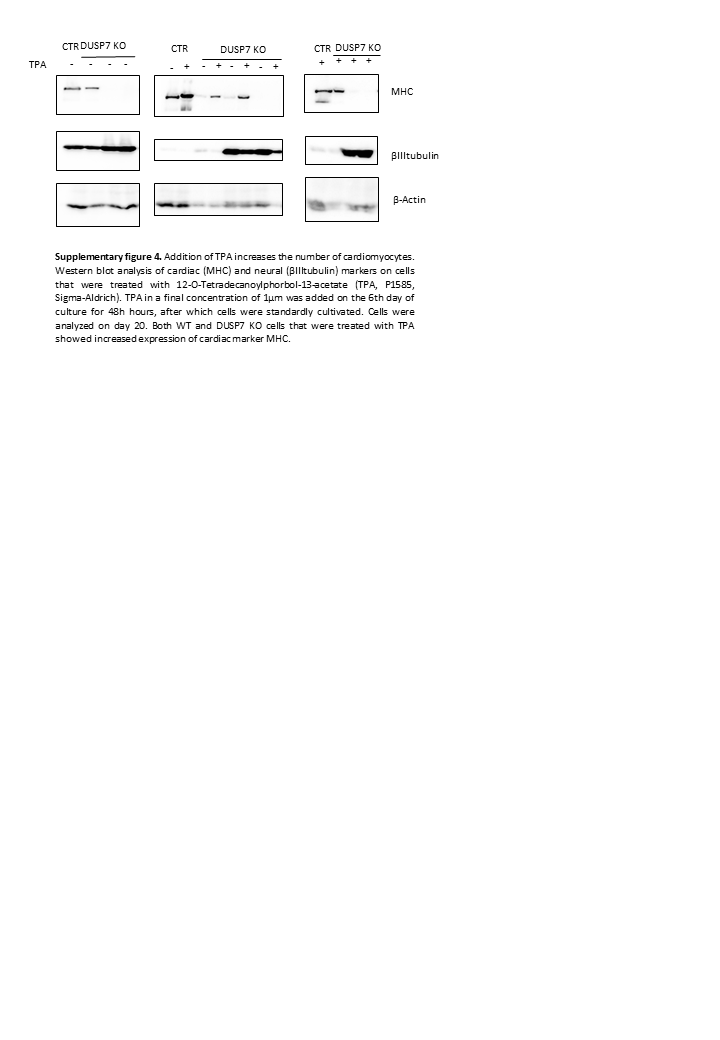

### Supplementary Figure 5

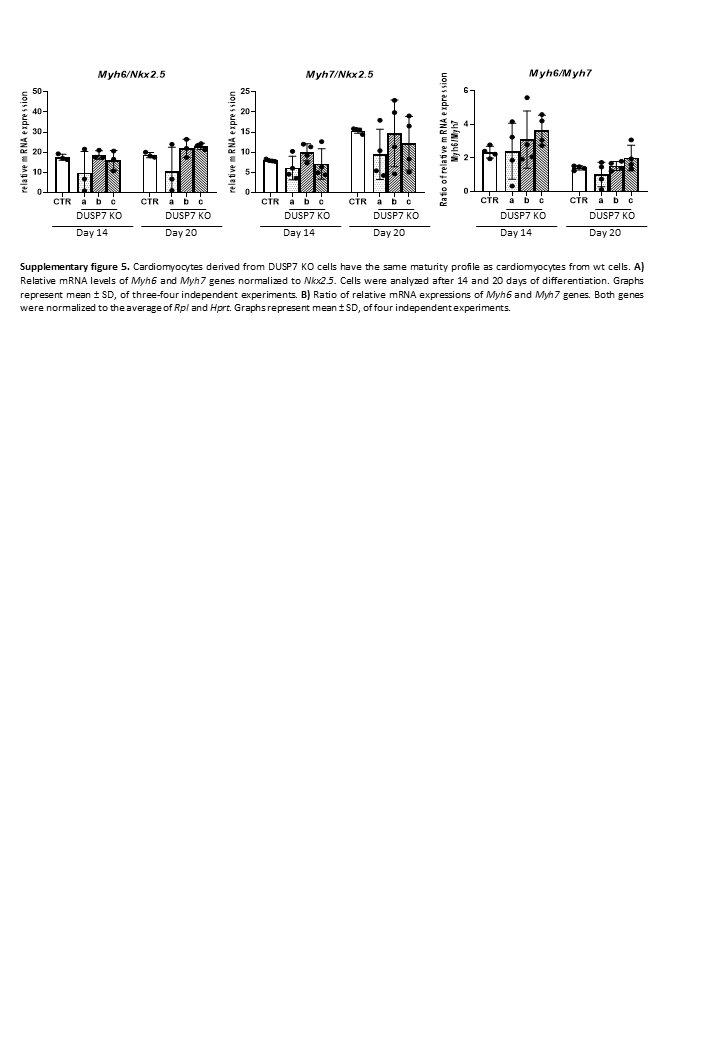

### Supplementary Figure 6

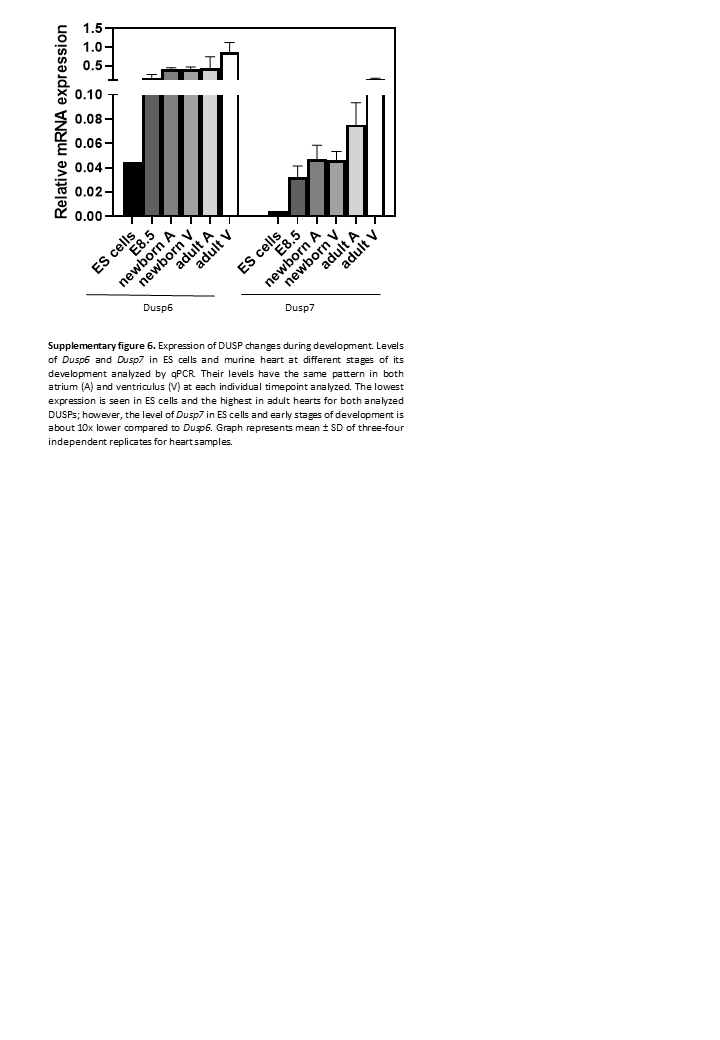

### Supplementary Figure 7

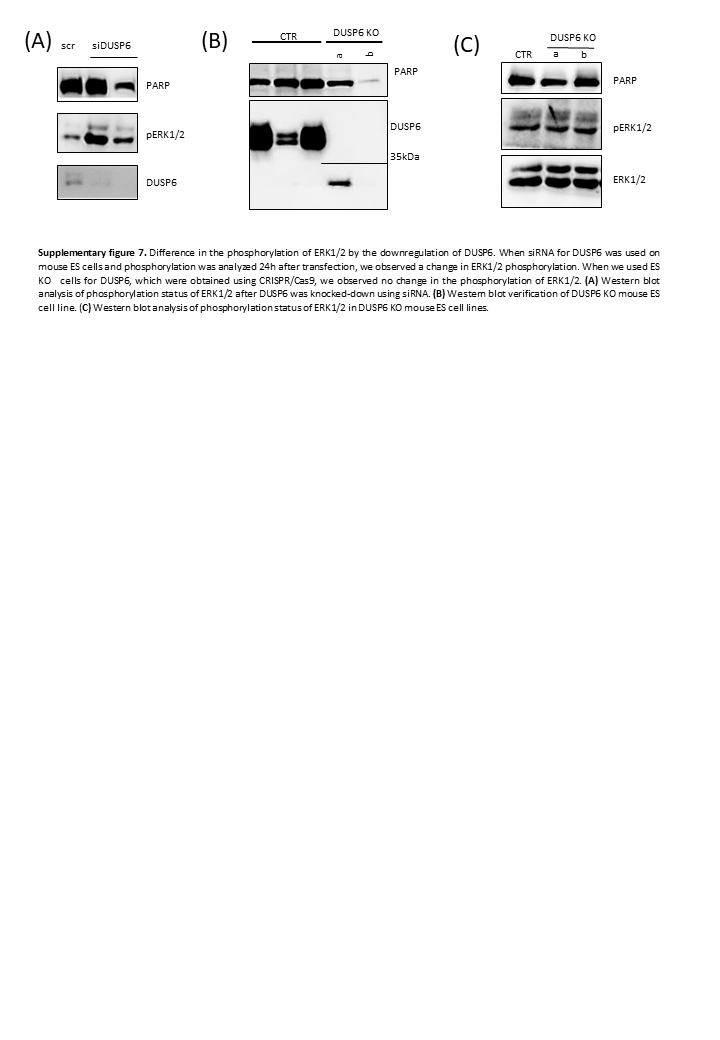
